# Targeting FGFR4 Abrogates HNF1A-driven Metastasis in Pancreatic Ductal Adenocarcinoma

**DOI:** 10.1101/2025.02.06.636643

**Authors:** Katherine J. Crawford, Kennedy S. Humphrey, Eduardo Cortes, Jianxin Wang, Michael E. Feigin, Agnieszka K. Witkiewicz, Erik S. Knudsen, Ethan V. Abel

**Affiliations:** Department of Molecular and Cellular Biology, Roswell Park Comprehensive Cancer Center, Buffalo, New York; Department of Pharmacology and Therapeutics, Roswell Park Comprehensive Cancer Center, Buffalo, New York; Department of Biostatistics and Bioinformatics, Roswell Park Comprehensive Cancer Center, Buffalo, New York; Department of Biostatistics, University at Buffalo, Buffalo, New York

**Keywords:** PDAC, pancreatic ductal adenocarcinoma, HNF1A, FGFR4, metastasis, FGFR4 inhibitors

## Abstract

**Purpose:** We previously identified an oncogenic role for the transcription factor HNF1A in pancreatic ductal adenocarcinoma (PDAC). However, the role of HNF1A in the metastatic progression of PDAC remains unknown and targeting modalities for HNF1A -dependent phenotypes have yet to be identified.

**Experimental Design:** Transwell chambers were used to assess the effects of HNF1A and FGFR4 modulation on the migration and invasion of ATCC and patient-derived PDAC cells *in vitro*. An intrasplenic injection xenograft model was used to evaluate the impact of HNF1A knockdown and overexpression on metastatic tumor burden. Single-cell RNA sequencing, tissue microarray (TMA) data, and UMAP spatial profiling were used to identify FGFR4 as an HNF1A target gene upregulated in metastatic cells. RNAi and two FGFR4 inhibiting modalities (H3B-6527 and U3- 1784) were utilized to demonstrate the efficacy of FGFR4 inhibiting agents at reducing HNF1A- driven metastasis.

**Results:** Knockdown of HNF1A significantly decreases and HNF1A overexpression significantly increases PDAC cell migration and invasion. *In vivo* studies show that HNF1A knockdown significantly abrogates metastasis, while overexpression significantly promotes metastasis. Single-cell RNAseq shows that FGFR4 is upregulated in metastatic PDAC cells and staining for HNF1A and FGFR4 in a PDAC TMA reveals significant correlation between HNF1A and FGFR4 in PDAC patients. Further, knockdown and inhibition of FGFR4 significantly decreases HNF1A- mediated cell migration and invasion, and blocks HNF1A-driven metastasis *in vivo*.

**Conclusions:** These findings demonstrate that HNF1A drives PDAC metastasis via upregulation of FGFR4, and FGFR4 inhibition is a potential mechanism to target metastasis in PDAC patients.

**Translational Relevance:** Pancreatic ductal adenocarcinoma (PDAC) is one of the most lethal malignancies, made even more devastating when metastases overwhelm major organs. The vast majority of PDAC patients either present with metastases or will relapse with recurrent metastatic PDAC after primary tumor resection. Unfortunately, toxic and largely ineffective chemotherapies are currently the only approved treatment options for these patients and therefore there exists a critical and unmet clinical need for targeted therapies against pro-metastatic pathways in PDAC. In the current study, we identify HNF1A as an oncogenic transcription factor that drives metastasis in PDAC, and it does so through upregulation of the receptor tyrosine kinase FGFR4. Importantly, FGFR4 is a targetable vulnerability and treatment with an FGFR4 blocking antibody reduces HNF1A-driven metastasis. These findings suggest that FGFR4 inhibitors could be an efficacious treatment for PDAC patients for the prevention or delay of metastatic tumor development.

## Introduction

Pancreatic ductal adenocarcinoma (PDAC) is the most common form of pancreatic cancer (∼90%) and is currently the 3^rd^ leading cause of cancer-related deaths, with a dismal 5-year survival rate of only 12.8%^1^. This poor prognosis is predominantly attributed to metastatic disease, as over 50% of patients present with metastatic disease, making them ineligible for curative surgery^1^. For these patients, who often succumb to the disease in less than one year, new treatment options are urgently needed. Further demonstrating the aggressiveness of PDAC metastases, approximately 80% of patients with clinically localized PDAC who undergo surgical resection will still die from recurrent metastatic PDAC^2^. As largely ineffective and toxic chemotherapeutics are the only treatment option available to metastatic and relapsed patients, there is a critical and unmet need for a better understanding of the pathways driving PDAC metastasis and identification of actionable drivers of these pathways.

Hepatocyte nuclear factor 1 α (HNF1A) is a gastrointestinal lineage transcription factor that controls the differentiation of liver and pancreatic cells^3–5^. We previously demonstrated an oncogenic role for HNF1A in PDAC. In functional studies, HNF1A overexpression transformed non-tumorigenic pancreatic cells and knockdown of HNF1A significantly depleted tumor growth in a patient-derived xenograft model^6^. Studies in prostate, renal, and colorectal cancers further support an oncogenic function for HNF1A^7–9^. Recent work by Cai *et al*. found that the HNF1A binding motif is the only transcription factor motif significantly enriched in the super enhancers of metastatic colorectal cancer and PDAC cell lines as compared to primary cell lines, suggesting a role for HNF1A in PDAC metastasis^10^. However, studies investigating the role of HNF1A in metastatic progression are limited.

Fibroblast growth factor receptor 4 (FGFR4) is a transcriptional target gene of HNF1A^6,11^. FGFR4 is a transmembrane receptor tyrosine kinase (RTK) that is normally expressed in the liver and regulates bile acid synthesis in response to digestion^12,13^. As with many RTKs, FGFR4 activation by ligand binding triggers phosphorylation and subsequent internal signaling cascades, including the PI3K/Akt and MAPK pathways, to induce cellular responses such as proliferation, differentiation, metabolism, and migration. FGFR4 serves an oncogenic function in several cancer types and promotes metastatic spread in other gastrointestinal cancers^14–19^.

Furthermore, FGFR4 is a driving oncogene in a subset of hepatocellular carcinomas in which FGF19, a unique ligand of FGFR4, is amplified. Because of this, several FGFR4-specific inhibitors have been developed and are currently undergoing clinical trials. H3B-6527, an FGFR4-specific tyrosine kinase inhibitor and U3-1784, an FGFR4 blocking antibody, were both utilized in this study^20,21^. Nevertheless, the function of FGFR4 in the metastatic progression of PDAC has yet to be elucidated.

In the current study, we aimed to determine whether HNF1A serves a role in promoting metastasis of PDAC, and the mechanisms by which it does so. Using *in vitro* and *in vivo* models, we identified HNF1A as a key driver of metastasis via transcriptional regulation of its target gene FGFR4. Knockdown of HNF1A significantly depleted metastatic tumors, and HNF1A overexpression significantly increased cell migration and invasion in several PDAC cell lines, which was abrogated by FGFR4 genetic knockdown and pharmacological inhibition. Most importantly, FGFR4 blockade was able to reduce HNF1A-driven metastasis in mice. These results suggest that FGFR4 inhibition is a promising avenue for delaying metastatic spread to extend survival in PDAC patients.

## Methods and Materials

### Cell culture

Human pancreatic cancer cell line AsPC-1 was purchased from ATCC (Manassas, VA). UM5 and UM53 (formerly NY5 and NY53, respectively) cell lines were established as low-passage cells isolated from human patient-derived xenografts as previously described^6^. All cell lines used were maintained in RPMI 1640 media supplemented with 10% FBS, 1% antibiotic-antimycotic, and 100ug/mL gentamicin. Cells were incubated at 37° with 5% CO2. Cells were routinely tested for mycoplasma contamination using the MycoScope PCR Detection kit (Gentlantis, San Diego, CA) and authenticated by STR profiling (Roswell Park Comprehensive Cancer Center Genomics Shared Resource).

### Lentiviruses

pENTR/D-TOPO/LacZ and pENTR/D-TOPO/HNF1A were generated as previously described. Human FGFR4 was amplified from UM5 cDNA with primers 5’-CAGGCTCCGCGGCCGCCCCCTTCACCATGCGGCTGCTGCTGGCCCTGTTGG-3’ and 5’-TGGGTCGGCGCGCCCACCCTTTCATGTCTGCACCCCAGACCCGAAGG-3’, digested with SacII and Ascl, and cloned into pENTR/D-TOPO using NEBuilder HiFi DNA Assembly master mix and protocol (New England Biolabs, Ipswich, MA). All cDNAs were shuttled into pLenti6.3/UbC/V5-DEST using LR Clonase II enzyme mix and protocol. miR30-based non-targeting control shRNA (5’-AGCGATCTCGCTTGGGCGAGAGTAAGTAGTGAAGCCACAGATGTACTTACTCTCGCCCAAGCGAGAG GGCACTCTCGCTTGGGCGAGAGTAAGTACATCTGTGGCTTCACTACTTACTCTCGCCCAAG CGAGAT -3’) and HNF1A targeting shRNA (5’-AGCGAGTCCCTTAGTGACAGTGTCTATAGTGAAGCCACAGATGTATAGACACTGTCACTAAGGGACCGGCAGGTCCCTTAGTGACAGTGTCTATACATCTGTGGCTTCACTATAGACACTGTCACTAAGGGACT-3’) were shuttled into pLentipuro5/TRE3G/V5-DEST, an all-in-one doxycycline inducible lentiviral vector developed using LR Clonase II enzyme mix and protocol. For labeling cells with firefly luciferase, PatGFP (a variant of EGFP containing the following mutations: S31R, Y40N, S73A, F100S, N106T, Y146F, N150K, M154T, V164A, I168T, I172V, A207V) was fused to the N-terminus of firefly luciferase Luc2 (Promega) (subcloned from pGL4.10) and cloned into pENTR/D-TOPO using Gibson 35 Assembly (New England Biolabs). PatGFP-Luc2 was recombined into pLenti0.3/EF/V5-DEST, a modified version of pLenti6.3/UbC/V5-DEST with the human EF-1α promoter instead of the human UbC promoter and no downstream promoter/selective marker cassette, to generate pLenti0.3/EF/GW/PatGFP-Luc2. To create the ZsGreen1 FGFR4 enhancer reporter, the multiple cloning site and minimal TA promoter from pTA-Luc (Clontech) was cloned upstream of ZsGreen1(Clontech) into pENTR/D-TOPO (Invitrogen, Waltham, MA) to create pENTR/D-TOPO/MCS-TA-ZsGreen1. The FGFR4 intronic enhancer region was amplified with the primers 5’-TCGATAGGTACCGCCGGCTGGAGCTGGGAGTGAGGCG-3’ and 5’-GAGTCTAGATCTGCCGGCGCGAAGACAGCCGCAGGGAC-3’, digested with KpnI and BglII, and subcloned into pENTR/D-TOPO/MCS-TA-ZsGreen1 to generate the wild-type entry vector. The HNF1A consensus site was mutagenized with the primers 5’-GGC<u>AAATTTGCGCGAAA</u>CCGCAGTGCACACAGGGCCTTTTG-3’ and 5’-CGG<u>TTTCGCGCAAATTT</u>GCCCCCTCCACCCCCTGCCGC-3’ to generate the HNF1A binding-defective mutant (underlined section is the HNF1A consensus site). Both the wild-type and mutant reporter cassettes were recombined into pLentineo3/BLOCK-iT-DEST with LR Clonase II (Invitrogen) to generate the reporter lentivirus constructs^6^. All viruses were packaged in 293FT cells. Cells were treated with lentivirus-containing conditioned media for 48-72 hours and then selected with blasticidin (cDNAs), puromycin (shRNAs), or G418 (ZsGreen1 reporter) for up to two weeks before use in studies. Pooled populations of transduced cells were continuously regenerated to prevent genetic drift.

### Inhibitors

H3B-6527 and U3-1784 for *in vitro* use were purchased from Selleck Chemicals (Houston, TX) and U3-1784 for *in vivo* use was purchased from Med Chem Express (Monmouth Junction, NJ). H3B-6527 was used at a concentration of 1 uM for the duration described in each figure. U3-1784 was used at a concentration of 5 ug/mL *in vitro* for the time indicated in each figure.

### siRNA knockdown

siRNAs were purchased from Dharmacon (Lafayette, CO) for Control (Cat# D-001810-01-20), HNF1A (Cat#M-008215-01-0005), FGFR4 sequence 10 (Cat #D—003134-10-0005), or FGFR4 sequence 11 (Cat #D-003134-11-0005). siRNA was combined with RNAiMax Lipofectamine (Invitrogen) in OptiMEM media and incubated at room temperature for 25 minutes before transfection. Knockdowns were confirmed via Western blot. siRNAswere transfected for the times indicated in each figure legend.

### Transwell assays

Cells were plated at a density of 1x10^5^ in 100 uL serum free media in the top chamber of Corning transwell 8 um inserts, or Corning Matrigel coated inserts for invasion assays, with 600 uL complete media plated in the bottom compartment. Cells were then incubated at 37° for 24 (AsPC-1 and UM5) or 6 (UM53) hours for migration assays and 48 (AsPC-1 and UM5) or 24 (UM53) hours for invasion assays. At endpoint, any cells remaining in the top chamber were removed with a cotton swab and cells on the reverse side of the chamber were fixed in 37% formaldehyde for 10 mins, dried for 10 mins, stained with crystal violet for 10 mins, and washed with water. Migrated/invaded cells were then counted. For migration and invasion assays testing FGFR4 inhibition or knockdown, cells were pretreated with drug or siRNA for 48 hours. Cells pretreated with drug were plated in serum free media containing drug.

### Western blotting

All lysates were collected and boiled in 1x Laemmli sample buffer with 2% β-mercaptoethanol for 5 minutes followed by electrophoresis on 4-20% Mini-PROTEAN TGX precast Tris-Glycine-SDS gels (Bio-Rad, Hercules, CA), transfer to low-fluorescent PVDF (Bio-Rad) and incubated overnight in primary antibodies (1 µg/ml) in 1x Animal-Free Blocking Solution (Cell Signaling Technology, Danvers, MA) plus 0.1% Tween-20. Blots were incubated in DyLight™ 700 or 800-conjugated secondary antibodies in 5% milk in TBS plus 0.1% Tween-20 and 0.005% SDS at room temperature for 1 hour and imaged/quantitated by an Odyssey® CLx imaging system (Li-Cor, Lincoln, NE). Antibodies were purchased from Cell Signaling Technology (FGFR4, HNF1A), Origene (ZsGreen), and Thermo Fisher Scientific (Actin).

### Intrasplenic injection

NOD/SCID/IL2γR^-/-^ (NSG) mice were bred and maintained at Roswell Park Comprehensive Cancer Center’s animal care facilities. Evenly mixed sex 14-15 week old mice were implanted with 500,000 AsPC-1 tumor cells in a volume of 100 µl of sterile PBS in the spleen of the mice. After a period of 5 minutes, the spleen was removed, and the splenic vessels cauterized.

Bioluminescent imaging was then performed at day 0, and once a week after that until endpoint to monitor tumor burden. For knockdown experiments, mice were put on a diet of doxycycline (2mg/ml)/5% sucrose water and doxycycline chow (VWR, Radnor, PA) to induce the expression of the non-targeting or HNF1A-targeting shRNAs starting one week prior to inoculation until endpoint. At collection, livers were divided into multiple pieces and fixed at different orientations to assure representative sections when cutting. For antibody experiments, mice were treated twice a week with 25 mg/kg via IP injection. Mice were sacrificed at the specified endpoint or if they exhibited signs of distress.

### Orthotopic injection

NOD/SCID/IL2γR^-/-^ (NSG) mice were bred and maintained at Roswell Park Comprehensive Cancer Center’s animal care facilities. Evenly mixed sex 7-8 week old mice were orthotopically implanted with 500,000 AsPC-1 tumor cells in a volume of 50 µl (1:1 volume of cell suspension in growth media and Matrigel) into the pancreas of the mice. Bioluminescent imaging was then performed at day 0, and once a week after that until endpoint to monitor tumor burden. For knockdown experiments, mice were put on doxycycline (2mg/ml)/5% sucrose water and doxycycline chow (VWR) to induce the expression of the non-targeting or HNF1A-targeting shRNAs. Mice were sacrificed at the specified endpoint or if they exhibited signs of distress.

### RNA-seq

The sequencing libraries were prepared from 200ng total RNA purified using the TruSeq Stranded Total RNA kit (Illumina Inc, San Diego, CA). Following manufacturer’s instructions, the first step depleted rRNA from total RNA. After ribosomal depletion, the remaining RNA was purified, fragmented and primed for cDNA synthesis. Fragmented RNA was then reverse transcribed into first strand cDNA using random primers. The next step removed the RNA template and synthesized a replacement strand, incorporating dUTP in place of dTTP to generate ds cDNA. AMPure XP beads (Beckman Coulter, Brea, CA) were used to separate the ds cDNA from the second strand reaction mix resulting in blunt-ended cDNA. A single ‘A’ nucleotide was then added to the 3’ ends of the blunt fragments. Multiple indexing adapters, containing a single ‘T’ nucleotide on the 3’ end of the adapter, were ligated to the ends of the ds cDNA, preparing them for hybridization onto a flow cell. Adapter-ligated libraries were amplified by PCR, purified using Ampure XP beads, and validated for appropriate size on a 4200 TapeStation D1000 Screentape (Agilent Technologies Inc., Santa Clara, CA). The DNA libraries were quantitated using KAPA Biosystems qPCR kit, and were pooled together in an equimolar fashion, following experimental design criteria. Each pool was denatured and diluted to 16 pM for On-Board Cluster Generation and sequencing on a HiSeq2500 sequencer using the appropriate paired-end cluster kit and rapid mode SBS reagents following the manufacturer’s recommended protocol (Illumina Inc.).

### TMA and multispectral staining

Multispectral immunofluorescent (mIF) staining was performed on formalin-fixed paraffin-embedded (FFPE) PDAC tissue microarray (TMA) using the Opal 6-Plex Detection Kit (AKOYA Biosciences, Marlborough, MA). FFPE samples were sectioned at 4µm thickness and placed on charged slides. The prepared slides were dried at 65°C for at least 2 hours. The dried slides were loaded into a BOND RX^m^ Research Stainer (Leica Biosystems) and deparaffinized with BOND Dewax solution (AR9222, Lecia Biosystems). The slides were treated with the Akoya Biosciences mIF staining process using Opal reagents. The process involved serial applications of the following solutions for each biomarker: ER1 (citrate buffer pH 6, AR996, Leica Biosystems) or ER2 (Tris-EDTA buffer pH9, AR9640, Leica Biosystems) epitope retrieval solutions, blocking buffer (Akoya Biosciences), primary antibody, PowerVision Poly-HRP (Leica Biosystems) secondary antibody, Opal fluorophore (Akoya Biosciences). Spectral DAPI (Akoya Biosciences #FP1490) was manually applied following removal of the samples from the BOND RX^m^. Processed slides were preserved with glass coverslips using ProLong Diamond Antifade Mountant (ThermoFisher Scientific). Antibodies used were purchased from Invitrogen (HNF1A, GT4110), Santa Cruz Biotechnology (FGFR4, A-10) and Agilent Dako (AE1/AE3). Slides were imaged on the PhenoImager HT Automated Quantitative Pathology Imaging System (AKOYA Biosciences). Further analysis of the slides was performed using in Form Software v2.6.0 225 (AKOYA Biosciences).

PDAC tumor cases were obtained from the surgical pathology files at Thomas Jefferson University. The TMA was contained specimens derived from largely consecutive cases between the years 2002 and 2010 under an Institutional Review Board approved protocol^22,23^. 220 PDAC patient cores with sufficient live cells (threshold > 100 cells per core) were stained and quantitated. Phenoptr software was used to report the percentage of cells that were positive for each protein stained. For correlation of staining intensities, average whole cell staining intensity was measured for each core. For the violin plot, the Pearson correlation coefficients were calculated for each core first. Then the R values from each core are used to make the violinplot. Likely dead cells were excluded from both percent positivity and staining intensity analyses.

### Immunohistochemistry

Anti-HNF1A antibody was purchased from Thermo Fisher Scientific (GT4110) and anti-Ki67 was purchased from Abcam (Cat #ab15580). Sections from FFPE tissues from the above mouse experiments were cut and stained with the above antibodies. Staining intensity was scored using 6 representative regions from each slide and calculated as the percent of positive cells.

### AlamarBlue cell viability assays

After the specified amount of time, cells were incubated in complete RPMI media containing 10% AlamarBlue reagent (Invitrogen) for 30 minutes-1 hour until a colorimetric change was observed. 100 uL of AlamarBlue containing media from each well was then plated in technical triplicates in a 96-well clear bottom/black-walled plate. Fluorescence was then measured on a BioTek Synergy HTX multi-mode reader on Gen5 3.03 software using 540nM excitation and 590nM emission wavelengths. All fluorescence was normalized to their respective control group set to 100% cell viability.

### Colony formation assays

200 cells were plated into 12-well plates incubated at 37° with 5% CO2 for 14 days. Media was replenished twice a week. At endpoint, cell viability was measured using AlamarBlue as outlined above followed by fixation with 5% formaldehyde and Crystal Violet staining for representative images. For knockdown studies, cells were transfected with siRNA 72 hours prior to plating for colony formation.

### Bioluminescent imaging

Bioluminescent imaging was performed on the IVIS Spectrum imager (Roswell Park Comprehensive Cancer Center Translational Imaging Shared Resource). Mice were anesthetized using isoflurane and injected intraperitoneally with 100uL of 15mg/mL luciferin. After a period of 5 minutes, mice were imaged from a supine position. Bioluminescence was quantified using the LivingImage software.

### Statistical analysis

Data throughout are expressed as the mean ± SEM. Statistically significant differences between two groups was determined by the two-sided Student t-test for continuous data, while ANOVA was used for comparisons among multiple groups with Tukey’s or Dunnett’s post hoc tests where appropriate, with significance defined as p < 0.05. GraphPad Prism 10 was used for these analyses. ns = non-significant, *p<0.05, **p<0.01, ***p<0.001, ****p<0.0001.

### Study approval

All animal protocols were approved by The Institute Animal Care and Use Committee (IACUC) at Roswell Park Comprehensive Cancer Center. The animal welfare assurance number for this study is A3143-01.

## Results

### HNF1A promotes PDAC cell migration and invasion

Based on its oncogenic role in PDAC^6^ and preliminary evidence as a metastatic driver^10^, we first aimed to establish a pro-metastatic function for HNF1A. To determine if HNF1A regulates PDAC metastasis, we first measured cell migration and invasion, as these are key characteristics of metastatic cells. We used transwell chambers to measure cells’ ability to migrate and invade after HNF1A modulation. siRNA mediated knockdown of HNF1A in two PDAC cell lines, AsPC-1 and UM5, which both endogenously express HNF1A (Figure 1A) resulted in significantly reduced PDAC cell migration through the transwell membrane as compared to cells transfected with a non-targeting control siRNA (Figure 1B). Cell invasion was measured as a cell’s ability to move through Matrigel coated transwell membranes, which recapitulate the extracellular matrix cells encounter when disseminating from the primary tumor. Consistently, cell invasion was also significantly decreased with HNF1A knockdown (Figure 1C).

**Figure 1.**
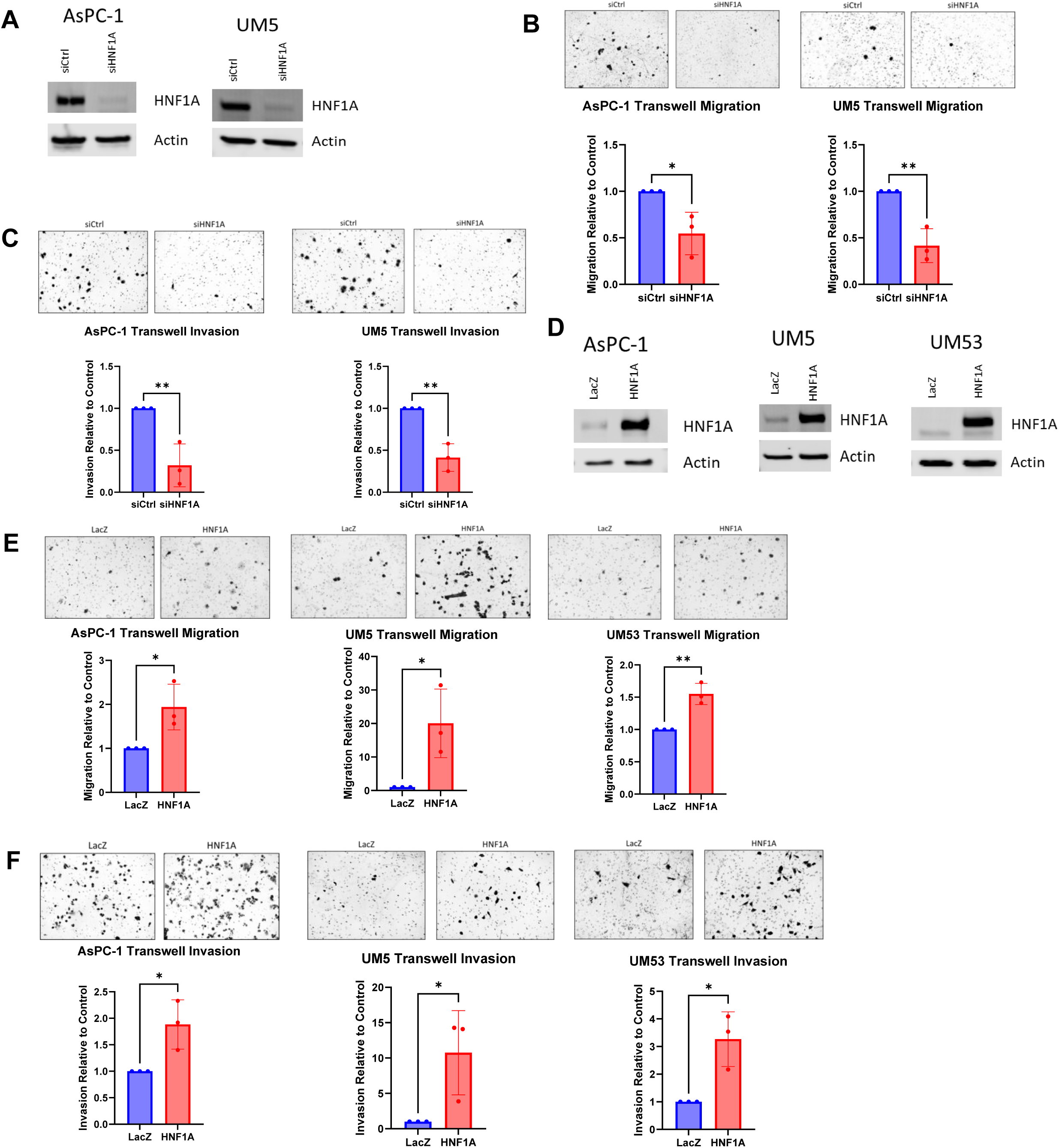
HNF1A promotes PDAC cell migration and invasion. A) Western blotting for HNF1A in AsPC-1 and UM5 cell lines to confirm knockdown. B) Normalized quantification and representative images of cell migration with HNF1A knockdown (n=3 biological replicates). Control knockdown or HNF1A knockdown cells were plated in transwell migration chambers and cells that had migrated after 24 hours were fixed and stained for counting. C) Normalized quantification and representative images of cell invasion with HNF1A knockdown (n=3 biological replicates). Control knockdown or HNF1A knockdown cells were plated in transwell invasion chambers and cells that had invaded after 48 hours were fixed and stained for counting. D) Western blotting for HNF1A in AsPC-1, UM5, and UM53 cells lines to confirm overexpression of HNF1A. E) Normalized quantification and representative images of cell migration with HNF1A overexpression (n=3 biological replicates). LacZ or HNF1A overexpressing cells were plated in transwell migration chambers and cells that had migrated after 6-24 hours were fixed and stained for counting. F) Normalized quantification and representative images of cell invasion with HNF1A overexpression (n=3 biological replicates). LacZ or HNF1A overexpressing cells were plated in transwell invasion chambers and cells that had migrated after 24-48 hours were fixed and stained for counting. All bar graphs represent the mean ± SEM and statistical difference was determined by unpaired t-test.

To evaluate whether HNF1A expression is sufficient to drive migration and invasion, we used lentiviral transduction to ectopically overexpress HNF1A in the UM53 cell line, which does not express HNF1A endogenously, in addition to AsPC-1 and UM5 cell lines. HNF1A overexpression in all three cell lines (Figure 1D) significantly increased both cell migration and invasion when compared to the LacZ expressing control cells. This effect was most dramatic in the UM5 cells, with over a 10-fold increase in both migration and invasion with HNF1A overexpression (Figure 1E and F). Together, these data demonstrate that HNF1A promotes PDAC cell migration and invasion *in vitro*.

### HNF1A drives PDAC liver metastasis in vivo

Because HNF1A promoted migratory and invasive capabilities, we next sought to determine if HNF1A drives PDAC liver metastasis *in vivo*. Initial efforts to establish spontaneous metastases with our cell models using orthotopic implantation resulted in exceedingly low penetrance of detectable metastases and circulating tumor cells (Supp Figure 1).

As such, we then moved on to use the intrasplenic injection liver seeding model of metastasis^24–26^, where shNTC or shHNF1A GFP-luciferase tagged AsPC-1 cells were injected intrasplenically into NSG mice (n=5). Bioluminescent imaging was then performed at day 0, then once a week for 4 weeks to assess the ability of the injected cells to seed and colonize the liver. At endpoint, mice were sacrificed, and the lung, liver, and pancreas from each mouse were harvested for formalin fixation and paraffin embedment (FFPE).

HNF1A depletion with doxycycline administration was confirmed via Western blot before inoculation and in tumors via IHC after harvest (Supp Figure 2A and B). Of note, HNF1A knockdown did not significantly impact cell viability, as both NTC and HNF1A knockdown injected cells produced bioluminescence, which can only be produced by living cells, at the Week 1 timepoint (data not shown). At the four-week endpoint, bioluminescent imaging showed a nearly complete abrogation of liver metastasis with knockdown of HNF1A (Figure 2A and B). FFPE liver samples were also H&E stained to view histological differences between metastases derived from shNTC or shHNF1A cells. Representative sections of livers were quantified measuring total metastatic tumor area as a percent of the whole liver area. HNF1A knockdown resulted in a near complete elimination of metastatic lesions, with a significant decrease in overall metastatic tumor area from 6.08% in the shNTC livers to 0.06% in the shHNF1A livers (Figure 2C and D). These data suggest that HNF1A expression is required for liver metastatic colonization.

**Figure 2.**
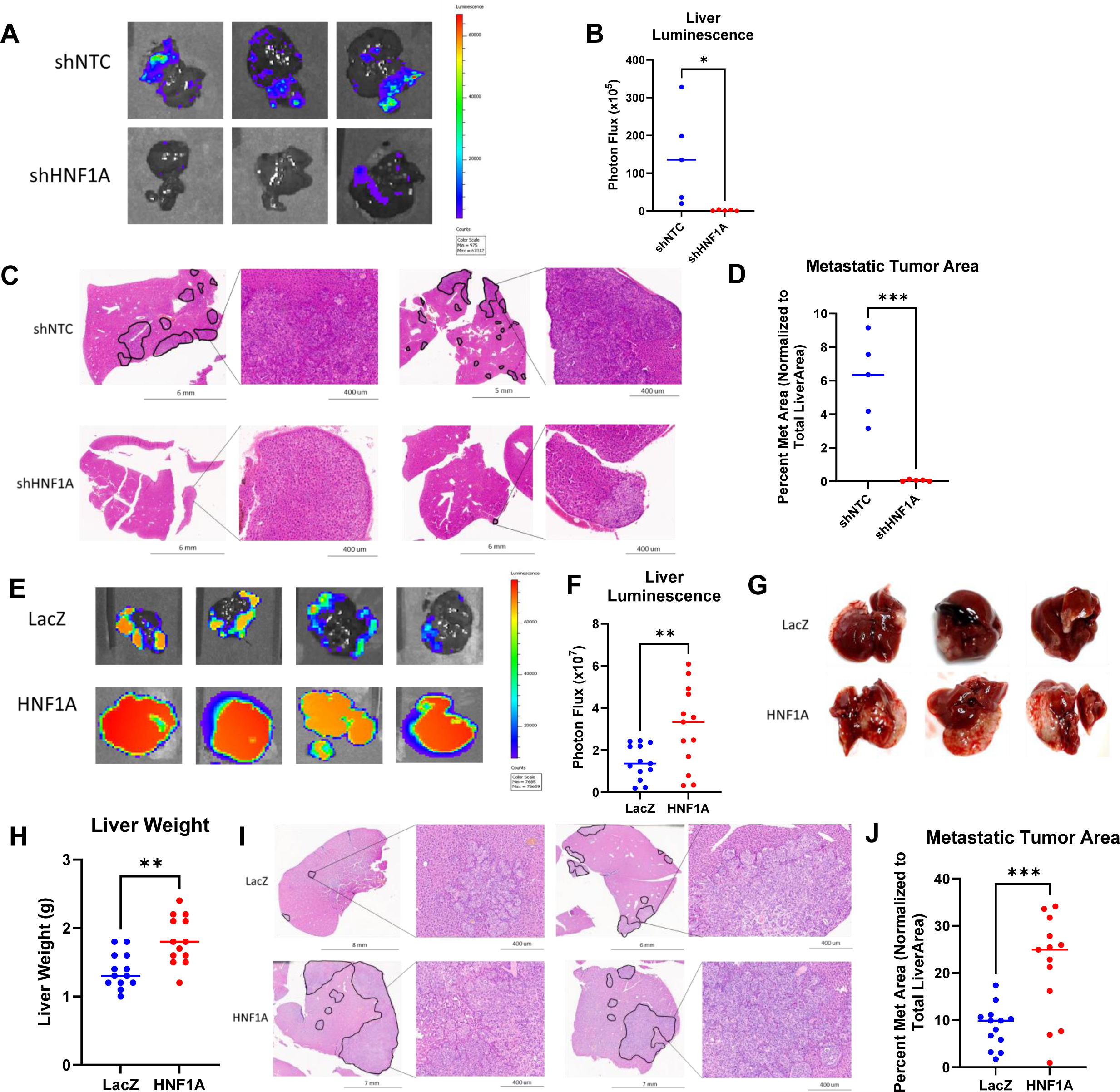
HNF1A drives PDAC liver metastasis *in vivo*. A) Representative bioluminescent images of harvested livers from mice implanted with control and HNF1A knockdown cells. B) Quantification of photon flux from bioluminescent images from all harvested livers at endpoint (n=5 mice per group). C) Representative images of H&E stained sections of liver tissue from both control and HNF1A knockdown groups. Black outlines indicate metastatic tissue. Zoomed in insets show histology of metastatic lesions. D) Quantification of total metastatic tumor area normalized as a percentage of total liver area (n=5 mice per group). E) Representative bioluminescent images of harvested livers from mice implanted with LacZ and HNF1A overexpressing cells. F) Quantification of photon flux from bioluminescent images from all harvested livers at endpoint (n=13 mice per group). G) Representative photographs of harvested livers from both LacZ and HNF1A groups showing increased tumor burden (white tissue) in the HNF1A livers. H) Measured liver weights of all livers after harvest at endpoint (n=13 mice per group). I) Representative images of H&E stained sections of liver tissue from both LacZ and HNF1A overexpression groups. Black outlines indicate metastatic tissue. Zoomed in insets show histology of metastatic lesions. J) Quantification of total metastatic tumor area normalized as a percentage of total liver area (n=13 mice per group). All bar graphs represent the mean and statistical difference was determined by unpaired t-test.

We next conducted the inverse experiment, in which the same intrasplenic injection was performed with GFP-luciferase tagged AsPC-1 cells overexpressing either LacZ control or HNF1A in NSG mice (n=13), to test whether HNF1A expression can induce PDAC metastasis. HNF1A overexpression was confirmed via Western blot before implantation of cells (Supp Figure 2C). Bioluminescent imaging was again performed at day 0 and once a week for 6 weeks. Imaging of the harvested livers at the six-week endpoint revealed significantly increased tumor burden in the livers of mice implanted with HNF1A overexpressing cells (Figure 2E and F). Images of the harvested livers also visibly show a difference in the extent of the metastatic lesions between those seeded with LacZ and HNF1A overexpressing cells (Figure 2G). The livers from HNF1A overexpressing mice also weighed significantly more than LacZ livers, again indicating increased tumor burden (Figure 2H). H&E staining of liver sections showed that HNF1A overexpression significantly increased the number and size of metastases, with a median of 8.62% of the total liver area being metastatic tissue in the control mice versus over 21% in the HNF1A livers (Figure 2I and J). Lung tissues were also collected from mice at endpoint and subjected to H&E staining to assess metastatic tumor development in other organs. While only 3 mice injected with LacZ overexpressing cells developed lung metastases, 8 mice that received HNF1A overexpressing cells developed lung metastases. HNF1A overexpression also resulted in more metastases per lung section (Supp Figure 2D).

As HNF1A overexpression has previously been shown to increase cell proliferation^6^, we further sought to validate that the metastatic tumors from mice inoculated with HNF1A overexpressing cells were not larger simply because they proliferated more rapidly. IHC staining was performed for the proliferative marker Ki67 on the slides from three representative mice from each group and found that the number of Ki67+ cells was not significantly different between LacZ and HNF1A overexpressing tumors (Supp Figure 2E). From these data, we concluded that HNF1A expression is necessary and sufficient to promote metastasis.

### HNF1A and FGFR4 expression correlate with human metastatic PDAC

As a transcription factor, HNF1A is currently not a druggable target, limiting its potential as a therapeutic vulnerability. As such, we aimed to identify a targetable downstream mediator of HNF1A-driven metastasis. Taking an unbiased approach, we analyzed publicly available single cell RNA-sequencing data (GSE 253260) to assess differentially expressed genes between metastatic tumors and primary tumors from PDAC patients. Of the significantly differentially expressed genes, 1,740 genes were upregulated in metastases vs primary tumors (log2 fold change > 1.5). We then compared this list of upregulated genes to genes significantly downregulated with HNF1A knockdown from our bulk RNA-sequencing in AsPC-1 cells. A substantial number of genes (252) overlapped between these two conditions suggesting that HNF1A may regulate numerous metastasis-associated genes. Of the genes that are both downregulated by HNF1A KD and upregulated in metastases, FGFR4 was a top hit (Figure 3A). FGFR4 is a known regulator of metastasis in other cancer types and can be therapeutically targeted^16,17,31,18–21,27–30^, though it’s role in PDAC metastasis is uncharacterized. Consistent with a role in PDAC metastasis, spatial FGFR4 expression from the single cell dataset in a UMAP plot showed significant overlap of FGFR4 expression with metastatic tumor cells as compared to primary tumor cells, indicating that FGFR4 expression is higher in metastatic cells as compared to primary tumor cells (Figure 3B).

**Figure 3.**
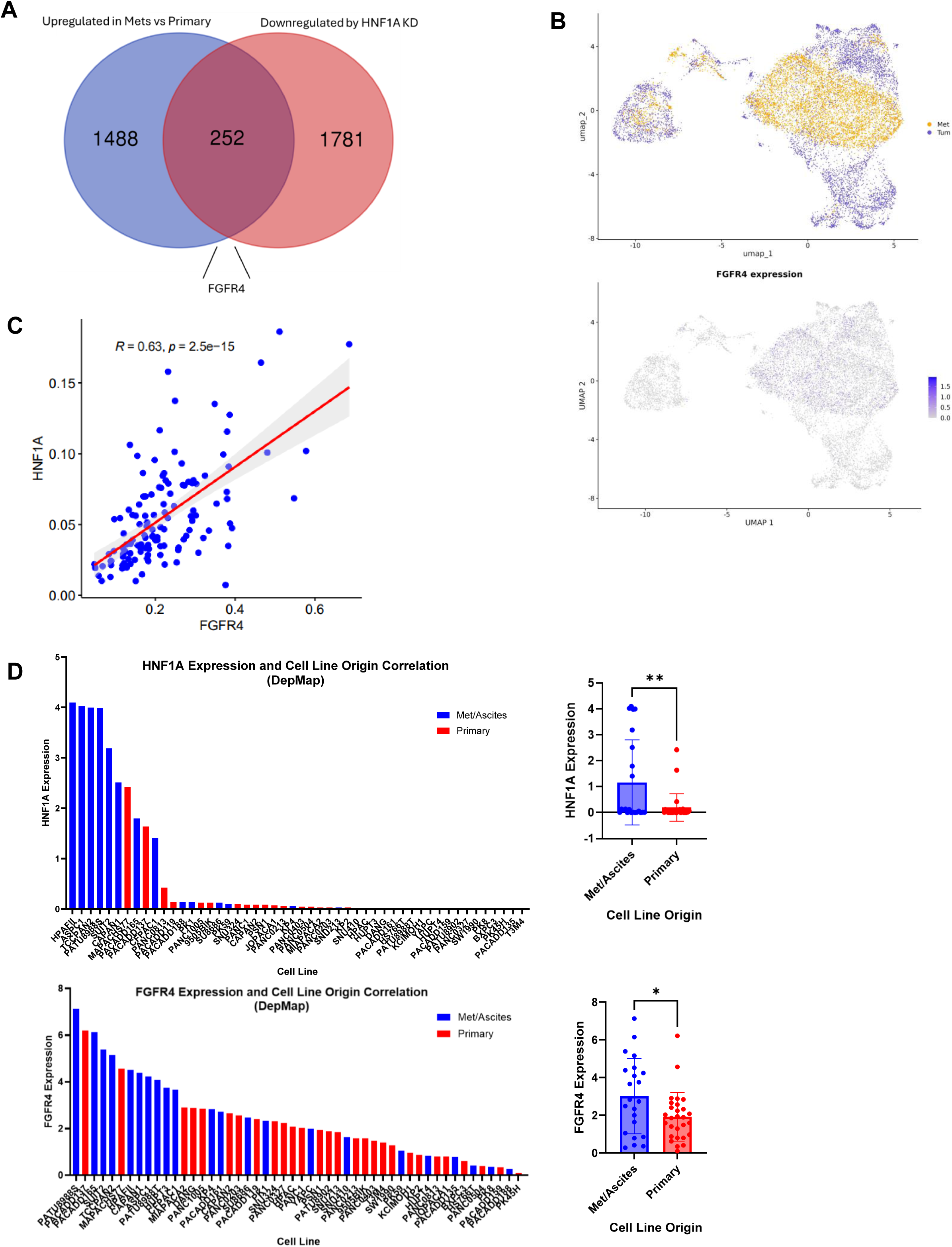
HNF1A and FGFR4 are associated with human metastatic PDAC. A) Venn diagram comparison of genes upregulated in metastatic tumors vs primary tumors from GSE253260 and genes significantly downregulated with HNF1A knockdown in AsPC-1 cells. B) Top: UMAP plotting of single cell RNA-sequencing showing spatial grouping of primary (purple) and metastatic (yellow) cells. Bottom: UMAP plotting FGFR4 expression in all single cells. C) Scatter plot comparing HNF1A and FGFR4 average staining intensity from multispectral immunofluorescent staining of PDAC TMA. D) Left: Plotting of all PDAC cell lines available in DepMap Portal based on either HNF1A (top) or FGFR4 (bottom) expression and whether the cell line is derived from primary tumors (red) or metastatic tumors/ascites (blue). Right: Comparison of primary vs metastasis derived cell lines based on the HNF1A (top) or FGFR4 (bottom) expression. Bar graphs represent the mean ± SEM and statistical difference was determined by unpaired t-test.

To determine if HNF1A and FGFR4 expression are widely associated in PDAC patients, we utilized multispectral immunofluorescence staining of a tumor microarray of 220 PDAC patient samples. These analyses revealed that HNF1A and FGFR4 were significantly and positively correlated in patient tumors, with a correlation coefficient of 0.63 (Figure 3C). We next sought to assess whether HNF1A and FGFR4 expression are increased in PDAC cells of metastatic origin compared to primary-derived cells. As there is a lack of paired primary/metastatic human PDAC tumor models, we utilized gene expression data from the DepMap Portal containing 22 primary- and 28 metastasis/ascites-derived cell lines. We found that expression of HNF1A and FGFR4 were significantly higher in PDAC cell lines of a metastatic/ascites origin versus primary tumor origin (Figure 3D). Based on these data, we hypothesize that HNF1A promotes metastasis via transcriptional regulation of FGFR4.

### HNF1A directly regulates FGFR4 expression

Based on the strong correlation between HNF1A and FGFR4 expression in metastatic disease, we next sought to validate FGFR4 as a direct target gene of HNF1A. Previous ChIP-sequencing from our group found an HNF1A binding peak, which contains the HNF1A binding sequence, in the FGFR4 enhancer region found in the first intron of the FGFR4 locus^6,11^. This putative enhancer was previously shown to interact with HNF1A in PDAC cells *in vitro*, with ectopic expression of HNF1A being able to induce FGFR4 in HNF1A-negative/FGFR4-negative Panc-1 cells^11^. As such, we sought to determine if endogenous HNF1A in fact regulates FGFR4 expression in PDAC cells. We found that siRNA-mediated knockdown of HNF1A reduced FGFR4 expression by more than 50% in both AsPC-1 and UM5 cells lines (Figure 4A). Lentiviral overexpression of HNF1A increased FGFR4 protein expression by more than 2-fold in all cell lines (Figure 4B).

**Figure 4.**
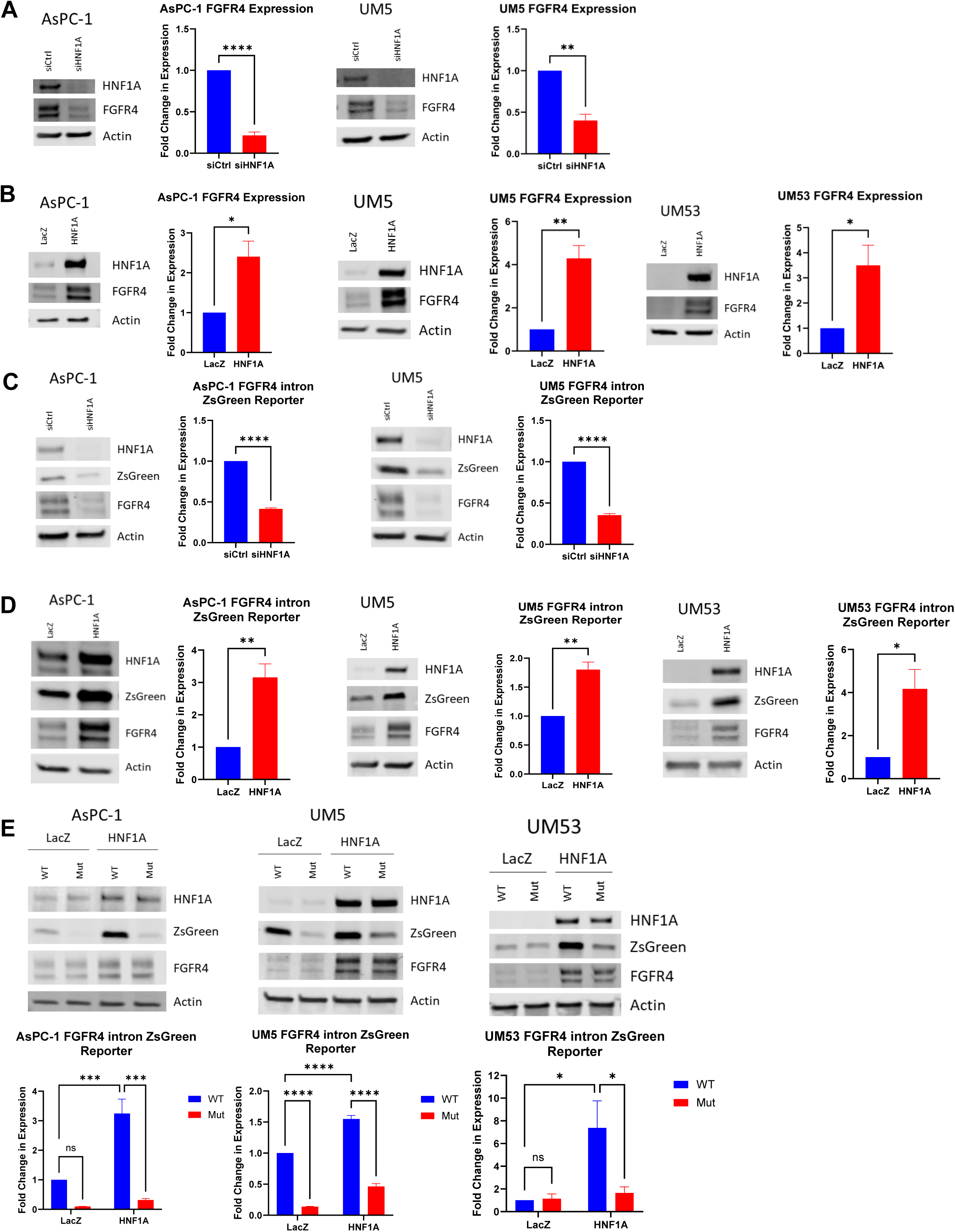
HNF1A directly regulates FGFR4 expression. A) Western blotting for HNF1A and FGFR4, and respective quantification of FGFR4 (n=3 biological replicates), with HNF1A knockdown in AsPC-1 and UM5 cell lines. B) Western blotting for HNF1A and FGFR4, and respective quantification of FGFR4 (n=3 biological replicates), with HNF1A overexpression in AsPC-1, UM5, and UM53 cell lines. C) Western blotting for HNF1A, FGFR4, and ZsGreen1 expression, and respective quantification of ZsGreen1 (n=3 biological replicates), in a reporter system with ZsGreen1 under the control of the FGFR4 enhancer with HNF1A knockdown in AsPC-1 and UM5 cell lines. D) Western blotting for HNF1A, FGFR4, and ZsGreen1 expression, and respective quantification of ZsGreen1 (n=3 biological replicates), in a reporter system with ZsGreen1 under the control of the FGFR4 enhancer with HNF1A overexpression in AsPC-1, UM5, and UM53 cell lines. E) Western blotting of HNF1A, FGFR4, and ZsGreen1, and respective quantification of ZsGreen1 (n=3 biological replicates), in the above reporter system with HNF1A overexpression +/- mutation of the HNF1A binding motif in the FGFR4 enhancer in AsPC-1, UM5, and UM53 cell lines. All bar graphs represent the mean ± SEM. Statistical difference was determined by unpaired t-test when comparing 2 conditions and one-way ANOVA with Tukey’s multiple comparisons test when comparing 4 conditions.

To confirm direct regulation of the FGFR4 enhancer region by HNF1A, we next designed a reporter in which ZsGreen1 expression is under the control of the FGFR4 enhancer sequence, which contains the HNF1A consensus sequence. Using this system, we found that HNF1A knockdown significantly depleted ZsGreen1 expression and that HNF1A overexpression significantly induced expression as compared to LacZ overexpressing control cells (Figure 4C and D). Finally, we mutated the HNF1A binding sequence in our reporter system, blocking the ability of HNF1A to bind the FGFR4 enhancer region. Mutation of this site resulted in a significant decrease in the HNF1A-induced ZsGreen1 expression (Figure 4E). Overall, we conclude that HNF1A directly binds an FGFR4 enhancer, promoting FGFR4 expression in PDAC.

### FGFR4 promotes pro-metastatic properties downstream of HNF1A

We next aimed to ascertain whether FGFR4, as a direct target gene of HNF1A, is responsible for the HNF1A-mediated metastatic phenotype. To evaluate this, we performed FGFR4 knockdown using two different siRNA sequences in both the LacZ and HNF1A overexpressing sublines of AsPC-1, UM5, and UM53 (Figure 5A). These cell line conditions were then plated for transwell migration and invasion assays, and we found that depletion of FGFR4 expression significantly reduced both the migration and invasion induced by HNF1A expression either back to or even below the LacZ siCtrl condition in all three cell lines (Figure 5B and C). Importantly, these effects were not due to a growth differential caused by knockdown of FGFR4. The two siRNA sequences did not significantly affect cell viability or colony formation in AsPC-1 or UM5 cells, though FGFR4 knockdown did have a significant impact on colony formation in UM53 cells (Supp Figure 3A and B).

**Figure 5.**
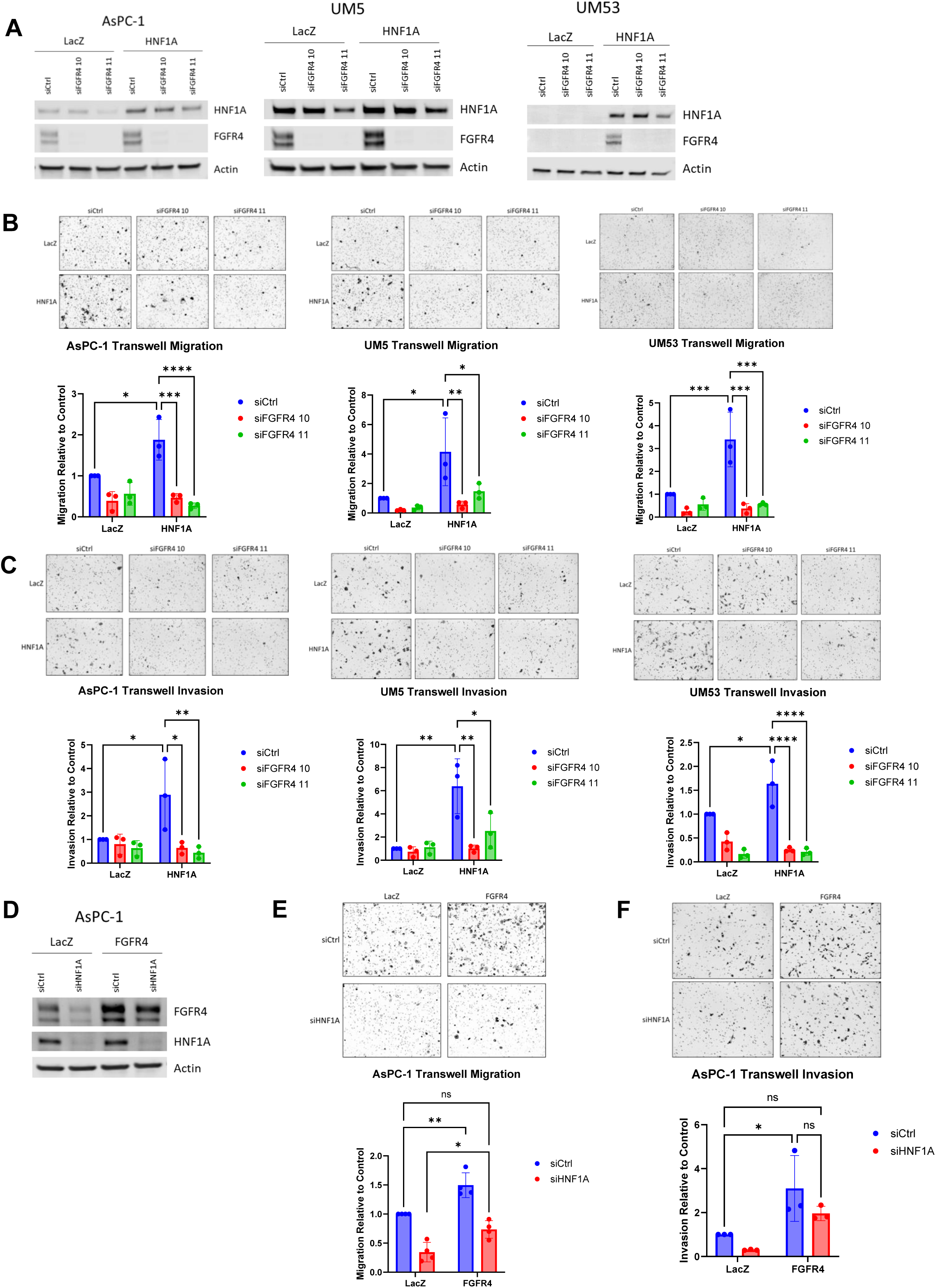
FGFR4 promotes migration and invasion downstream of HNF1A. A) Western blotting for HNF1A and FGFR4 to confirm HNF1A overexpression and FGFR4 knockdown in AsPC-1, UM5, and UM53 cell lines. B) Normalized quantification and representative images of cell migration in both LacZ and HNF1A overexpressing sublines with FGFR4 knockdown (n=3 biological replicates). Control or FGFR4 knockdown cells were plated in transwell migration chambers and cells that had migrated after 6-24 hours were fixed and stained for counting. C) Normalized quantification and representative images of cell invasion in both LacZ and HNF1A overexpressing sublines with FGFR4 knockdown (n=3 biological replicates). Control or FGFR4 knockdown cells were plated in transwell invasion chambers and cells that had migrated after 24-48 hours were fixed and stained for counting. D) Western blotting for HNF1A and FGFR4 to confirm FGFR4 overexpression and HNF1A knockdown in AsPC-1 cells. E) Normalized quantification and representative images of cell migration in both LacZ and FGFR4 overexpressing sublines with HNF1A knockdown (n=4 biological replicates). Control or FGFR4 knockdown cells were plated in transwell migration chambers and cells that had migrated after 24 hours were fixed and stained for counting. F) Normalized quantification and representative images of cell invasion in both LacZ and FGFR4 overexpressing sublines with HNF1A knockdown (n=3 biological replicates). Control or FGFR4 knockdown cells were plated in transwell invasion chambers and cells that had migrated after 48 hours were fixed and stained for counting. All bar graphs represent the mean ± SEM and statistical difference was determined by one-way ANOVA with Tukey’s multiple comparisons test.

To ascertain whether FGFR4 is sufficient to promote metastatic properties, we also tested whether FGFR4 overexpression could rescue cell migration and invasion from the decrease induced by HNF1A knockdown. FGFR4 overexpression and HNF1A knockdown in AsPC-1 cells (Figure 5D) revealed that FGFR4 overexpression alone significantly increased cell migration and invasion as compared to LacZ control cells. More importantly, overexpression of FGFR4 was able to rescue migration and invasion from the effects of HNF1A knockdown, as cells with HNF1A knockdown and FGFR4 overexpression migrated and invaded significantly more than those with HNF1A knockdown alone (Figure 5E and F). Again, the effects of FGFR4 are specific to migration and invasion as cell viability and colony formation were not significantly changed with FGFR4 overexpression, except for colony formation in UM5 cells (Supp Figure 2C and D). These data strongly support the hypothesis that FGFR4 is responsible for HNF1A-driven PDAC cell migration and invasion.

### Pharmacologic inhibition of FGFR4 ablates HNF1A-driven metastasis

We further wanted to elucidate whether pharmacologic inhibition of FGFR4 could recapitulate the effects of FGFR4 knockdown. FGFR4 targeting agents are an area of high interest in colorectal cancer, where a subset of patients exhibit an amplification of FGF19, a unique FGFR4 ligand. Because of this, several classes of FGFR4 inhibitors have been developed and tested in clinical trials. To test this the effects of pharmacologic blockade of FGFR4 on HNF1A-driven metastatic phenotypes, we used two FGFR4 inhibiting modalities; H3B-6527, an FGFR4-specific tyrosine kinase inhibitor^20^, and U3-1784, an FGFR4 blocking antibody^21^.

Consistent with the genetic manipulation findings, we found that treatment with H3B-6527 significantly decreased cell migration and invasion promoted by HNF1A overexpression in AsPC-1 cells (Figure 6A and B). Similarly, U3-1784 treatment was able to significantly reduce HNF1A-induced migration and invasion (Figure 6C and D). Treatment with these agents did not perturb total FGFR4 expression (Supp Fig 4). These results were not due to any effect on cell viability by the FGFR4 inhibitors, as cell viability and colony formation were only minimally impacted by treatment with either H3B-6527 or U3-1784 (Supp Fig 5).

**Figure 6.**
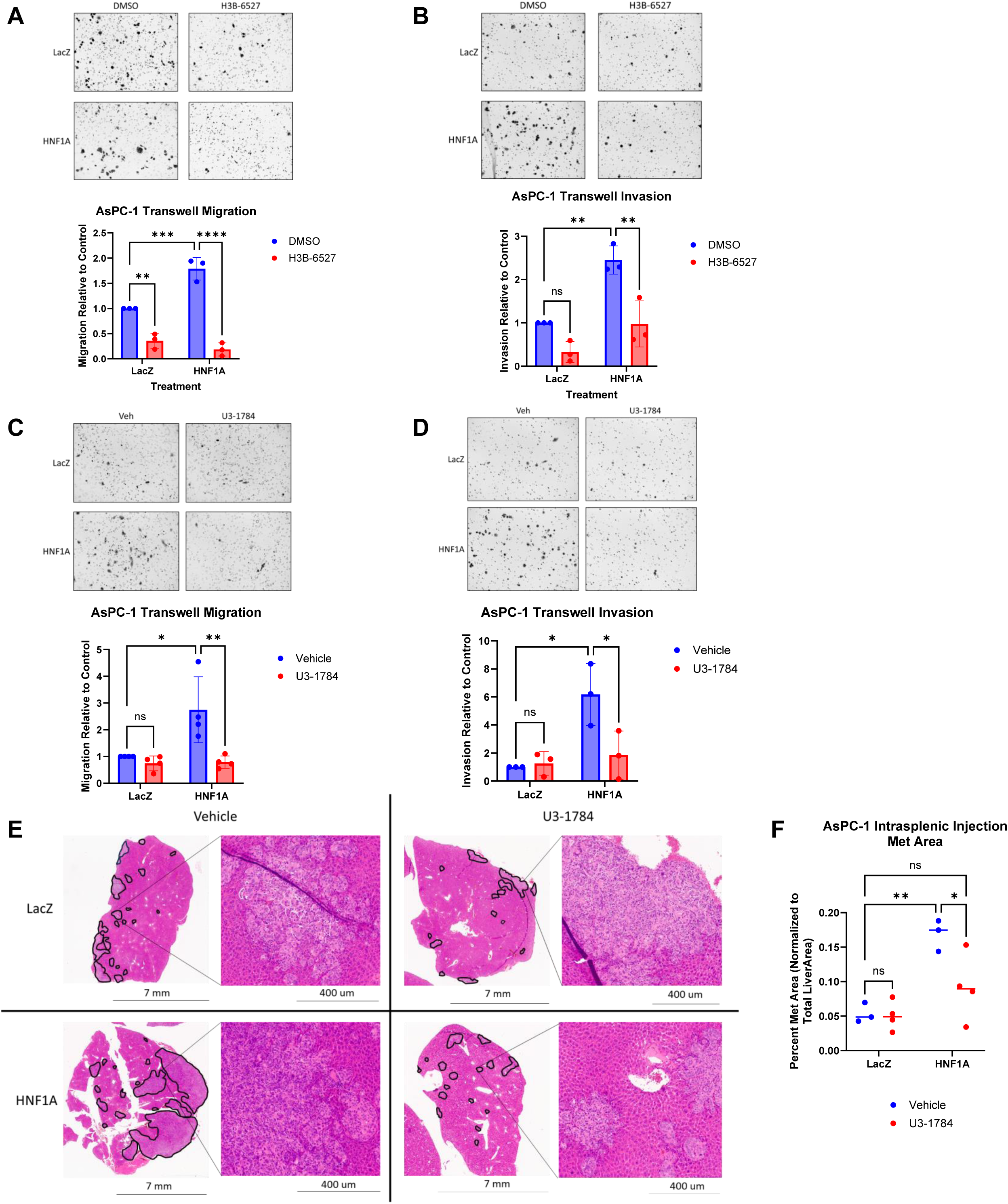
Pharmacologic inhibition of FGFR4 reduces HNF1A-driven metastasis. A) Normalized quantification and representative images of cell migration in both LacZ and HNF1A overexpressing sublines treated with 1μM H3B-6527 (n=3 biological replicates). DMSO or H3B-6527 treated cells were plated in transwell migration chambers in serum free media containing DMSO or H3B-6527 and cells that had migrated after 24 hours were fixed and stained for counting. B) Normalized quantification and representative images of cell invasion in both LacZ and HNF1A overexpressing sublines treated with 1μM H3B-6527 (n=3 biological replicates). DMSO or H3B-6527 treated cells were plated in transwell invasion chambers in serum free media containing DMSO or H3B-6527 and cells that had migrated after 48 hours were fixed and stained for counting. C) Normalized quantification and representative images of cell migration in both LacZ and HNF1A overexpressing sublines treated with 5μg/mL U3-1784 (n=4 biological replicates). Vehicle or U3-1784 treated cells were plated in transwell migration chambers in serum free media containing vehicle or U3-1784 and cells that had migrated after 24 hours were fixed and stained for counting. D) Normalized quantification and representative images of cell invasion in both LacZ and HNF1A overexpressing sublines treated with 5μg/mL U3-1784 (n=3 biological replicates). Vehicle or U3-1784 treated cells were plated in transwell invasion chambers in serum free media containing vehicle or U3-1784 and cells that had migrated after 48 hours were fixed and stained for counting. E) Representative images of H&E stained sections of liver tissue from both LacZ and HNF1A groups with either vehicle or U3-1784 treatment. Black outlines indicate metastatic tissue. Zoomed in insets show histology of metastatic lesions. F) Quantification of total metastatic tumor area normalized as a percentage of total liver area (n=3 mice per group for vehicle treated, 4 mice per group for U3-1784 treated). All bar graphs represent the mean ± SEM and statistical difference was determined by one-way ANOVA with Tukey’s multiple comparisons test.

Finally, to assess the efficacy of FGFR4 inhibition on reducing metastasis *in vivo*, we treated mice with either vehicle control or 25 mg/kg of U3-1784 twice a week for 4 weeks following intrasplenic injection of either LacZ or HNF1A overexpressing sublines of GFP-luciferase tagged AsPC-1 cells. Livers from these mice were harvested at endpoint and the FFPE tissues were H&E stained (Figure 6E). As before, total metastatic tumor area was measured and normalized to total liver area. Expectedly, the livers from mice inoculated with HNF1A overexpressing cells had significantly increased metastatic tumor burden are as compared to mice implanted with LacZ cells. However, treatment with the FGFR4 blocking agent significantly reduced the development and growth of metastatic lesions in the HNF1A livers (Figure 6F). Overall, these findings strongly indicate that treatment with an FGFR4 inhibitor is a clinically viable avenue for the disruption of HNF1A-driven metastasis in PDAC.

## Discussion

In the current study, we identified the transcription factor HNF1A as a driver of metastasis in PDAC, introducing a novel pathway in the progression of pancreatic cancer. Functional data reveal that loss of HNF1A dramatically reduces cell migration and invasion *in vitro*, and nearly eliminates PDAC liver metastasis *in vivo*. Our data further show that the receptor tyrosine kinase FGFR4 is a therapeutic vulnerability in this signaling axis. Knockdown or inhibition of FGFR4, using two different inhibitory agents, significantly reduced the HNF1A-mediated increase in migration, invasion, and metastasis.

One of the major takeaways from this work is the clinical applicability of FGFR4 inhibitors for the treatment of PDAC, and there are several treatment scenarios in which FGFR4 inhibiting agents are likely to be efficacious for PDAC patients. The first is as a neoadjuvant therapy for patients with resectable or borderline resectable primary tumors, categories which approximately 50% of PDAC patients fall into. Because the large majority of patients that undergo surgery will succumb to metastatic relapse^1,2^, FGFR4 inhibition before surgery could improve the potential for cure with resection by preventing recurrent metastatic tumors. Second, a small subpopulation of the public is regularly screened for pancreatic cancer due to their family history and/or risk factors. Similar to neoadjuvant treatment for surgery, treatment with FGFR4 inhibitors at the earliest signs of disease for these patients could prevent the characteristically early metastatic spread of PDAC, leading to an improved survival outcome.

Unfortunately, most PDAC patients present with distant metastases already present, leaving them with a bleak prognosis. Though FGFR4 inhibitors are unlikely to deplete existing metastatic tumors as they are not cytotoxic, preventing or delaying further lesions could allow standard of care chemotherapies to prove more efficacious and extend survival. Lastly, the work done in this study provides evidence that HNF1A and FGFR4 could serve as biomarkers for treatment stratification, as patients with HNF1A^high^ and/or FGFR4^high^ tumors will likely respond well to FGFR4 inhibitors. FGFR4 expression may also serve as a biomarker for response to erlotinib, while not a direct target itself, as recent work by Rao *et al.* showed that patients with HNF1A positivity had significantly better survival as compared to HNF1A negative patients when treated with Gemcitabine plus erlotinib^32^. Collectively, FGFR4 and responsiveness to erlotinib may indicate that HNF1A regulates a larger RTK network, which should be explored more extensively in future studies.

Other groups have previously identified tumor suppressive functions for both HNF1A and FGFR4^33,34^. However, in addition to our current findings demonstrating an oncogenic and pro-metastatic role for HNF1A, our group has previously published that HNF1A is an oncogene by promoting stem-like properties in PDAC cells^6^. Regarding FGFR4, one key divergence in our work from others is the selection of model systems. HPAF-II was the principal cell line model used in the study by D’Agosto *et al*.^33^. Our ability to replicate our findings across several cell lines, including primary PDAC cells, lends to the rigor of the results. We also observed consistent results using multiple modalities of FGFR4 modulation and inhibition, using both a tyrosine kinase inhibitor and blocking antibody. Moreover, the FGFR4 inhibiting therapies selected for use in this study are more clinically relevant, as they have been tested in humans, while BLU-9931 used by other groups never advanced to clinical trials due to the toxicity profile observed in mice^33,35^.

An encouraging finding of our work is the lack of cytotoxic effects with FGFR4 depletion or inhibition, lending to the study’s translatability. Unlike pan-FGFR inhibitors, or even some FGFR1-3 specific inhibitors, the two FGFR4 targeting modalities used in this work showed minimal effects on cell viability and colony formation, suggesting a lack of harsh toxicities with these therapies. Moreso, treatment with U3-1784 *in vivo* did not lead to any significant changes in body weight as compared to the vehicle treated groups, again indicating that this regimen is likely tolerable. Clinical trial data of other FGFR4 inhibitors found that one of the most common side effects of these agents is increased bile acid production, though this could be offset with the addition of a bile acid sequestrant^30,31,36^. Therefore, FGFR4 inhibition is a promising avenue for the safe and effective treatment of PDAC.

We acknowledge that a limitation of this work is the absence of a biomarker for the on-target effects of FGFR4 inhibition. Knockdown or inhibition of FGFR4 did not result in a change in any of the canonical FGFR4 downstream targets, including the PI3K/Akt and MAPK pathways, or in any metastasis-associated programs such as EMT (data not shown). Effects on migration, invasion, and metastasis served as functional readouts of FGFR4 loss, though a molecular biomarker was not found. Future work will therefore need to address this to identify a readout of the efficacy of the FGFR4 inhibitors used in this study, which will not only establish biomarkers of on-target inhibition but will also identify other potential vulnerabilities. One interesting finding from this study is the lack of effect of FGFR4 knockdown or inhibition in the LacZ context, even though these cells still express FGFR4. Though not tested in this study, it is possible that this is because of other HNF1A targets that may be involved in metastatic progression, independent of FGFR4. The comparison of HNF1A regulated genes and metastasis associated genes revealed that over 200 putative HNF1A target genes are correlated with metastasis. This suggests that there may be other players involved in HNF1A’s regulation of metastatic spread in PDAC and that these players are able to compensate for the loss of FGFR4 signaling. Direct inhibition of HNF1A expression and/or activity, such as via PROTACs, may therefore be a promising channel to overcome this challenge by blocking all HNF1A metastasis-promoting pathways.

In summary, we identified a novel role for HNF1A in the metastatic progression of PDAC through its transcriptional regulation of FGFR4. This therapeutic vulnerability may, in turn, be leveraged by using FGFR4 inhibitors to block or delay the spread of PDAC to vital organs and extend patient survival.

## Acknowledgements

We thank all the members of the laboratory group and colleagues in the discussion and preparation of this manuscript, especially the labs of Drs. Michael Feigin (RPCCC) and Anna Bianchi-Smiraglia (RPCCC). We would like to thank Sidney Mahan (RPCCC) and Hanna Rosenheck (RPCCC) for helping to optimize and perform multispectral immunofluorescence, and Robert Kyne (RPCCC) for assisting with animal studies. We would like to acknowledge Bryan Gillard (RPCCC) and Dr. Minhyung Kim (RPCCC) for performing all orthotopic and intrasplenic inoculation procedures. This work was supported by Roswell Park Comprehensive Cancer Center and National Cancer Institute (NCI) grant, P30CA016056; NCI grant R37CA275961; Pancreatic Cancer Action Network-AACR Pathway to Leadership Grant 16-70-25-ABEL; the SAS Foundation for Cancer Research Grant; the Hirshberg Foundation for Pancreatic Cancer Research Seed Grant (to EVA); and NCI grants CA267467 and CA211878 (to EK and AW). The content is solely the responsibility of the authors and does not necessarily represent the official view of the National Institutes of Health.

## Data Availability Statement

Raw data were generated in a core facility but processed data are available from the authors.

## Conflicts of Interest

The authors declare no potential conflicts of interest.

---

Supp Figure 1. A) Quantification of photon flux from bioluminescent images from all harvested livers and lungs inoculated with LacZ or HNF1A overexpressing cells at endpoint (n=10 mice per group). B) Quantification of photon flux from bioluminescent images from all harvested livers and lungs with control or HNF1A knockdown at endpoint (n=10 mice per group). C) Percent of blood cells positive for GFP (tumor cells) collected from the cardiac blood. All bar graphs represent the mean and statistical difference was determined by unpaired t-test.

Supp Figure 2. A) Western blotting for HNF1A with and without doxycycline administration before cell inoculation. B) Immunohistochemistry for HNF1A in the liver metastases of mice from each respective arm. C) Western blotting for HNF1A in AsPC-1 cells before inoculation. D) Quantification of the number of lung metastases per H&E stained tissue section. E) Quantification of cells positive for Ki67 immunohistochemistry staining in the liver metastases from 3 representative mice from each group. All bar graphs represent the mean and statistical difference was determined by unpaired t-test.

Supp Figure 3. A) Cell viability as normalized to control knockdown 72 hours after siRNA-mediated knockdown of FGFR4. B) Cell viability as normalized to control knockdown for colony formation experiments. Cells were transfected with siRNA for 72 hours before plating at a low cell density (200 cells/well) and grown for 2 weeks. Resultant colonies were quantified by AlamarBlue viability assay. C) Cell viability as normalized to LacZ control overexpression 72 hours after plating. D) Cell viability as normalized to LacZ control overexpression for colony formation experiments. Cells were plated at a low cell density (200 cells/well) and grown for 2 weeks. Resultant colonies were quantified by AlamarBlue viability assay. All bar graphs represent the mean and statistical difference was determined by unpaired t-test.

Supp Figure 4. A) Western blotting of HNF1A and total FGFR4 in AsPC-1, UM5, and UM53 cell lines treated with either DMSO or H3B-6527. B) Western blotting of HNF1A and total FGFR4 in AsPC-1, UM5, and UM53 cell lines treated with either Vehicle or U3-1784.

Supp Figure 5. A) Cell viability of LacZ and HNF1A overexpressing AsPC-1, UM5, and UM53 cells treated with DMSO or H3B-6527 as normalized to LacZ control cells treated with DMSO after 72-hour treatment. B) Cell viability for colony formation experiments as normalized to LacZ cells treated with DMSO. Cells were treated with wither DMSO or H3B-6527 for 72 hours before plating at a low cell density (200 cells/well) and grown for 2 weeks. Resultant colonies were quantified by AlamarBlue viability assay. C) Cell viability of LacZ and HNF1A overexpressing AsPC-1, UM5, and UM53 cells treated with Vehicle or U3-1784 as normalized to LacZ control cells treated with DMSO after 72-hour treatment. D) Cell viability for colony formation experiments as normalized to LacZ cells treated with DMSO. Cells were treated with wither DMSO or H3B-6527 for 72 hours before plating at a low cell density (200 cells/well) and grown for 2 weeks. Resultant colonies were quantified by AlamarBlue viability assay. All bar graphs represent the mean and statistical difference was determined by one-way ANOVA with Tukey’s multiple comparisons test.

## Notes

### Competing Interest Statement

The authors have declared no competing interest.

