## Supplementary figures and images for "Targeting FGFR4 Abrogates HNF1A-driven Metastasis in Pancreatic Ductal Adenocarcinoma"

### Supplemental Figures

Supp Figure 1

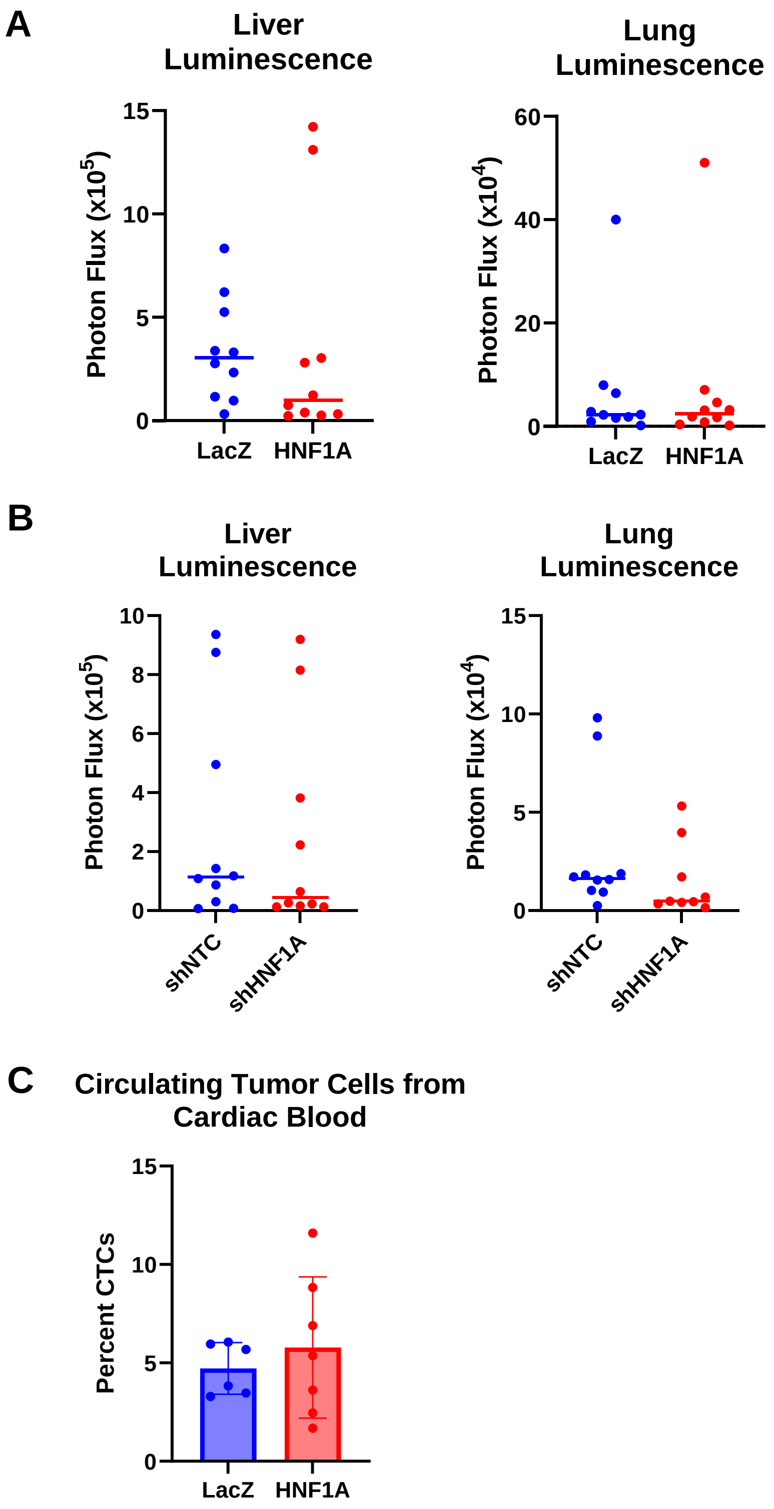

Supp Figure 2

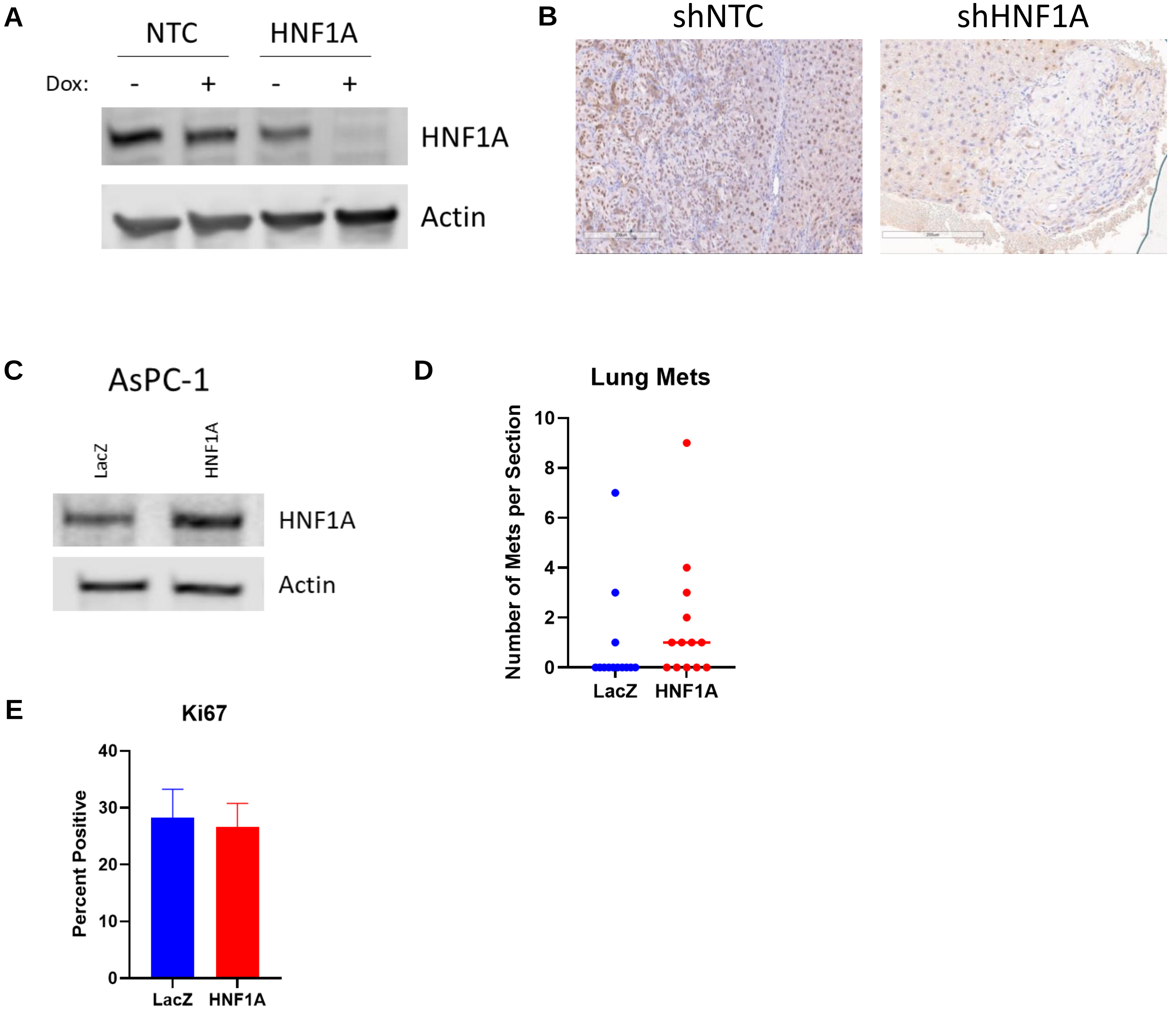

Supp Figure 3

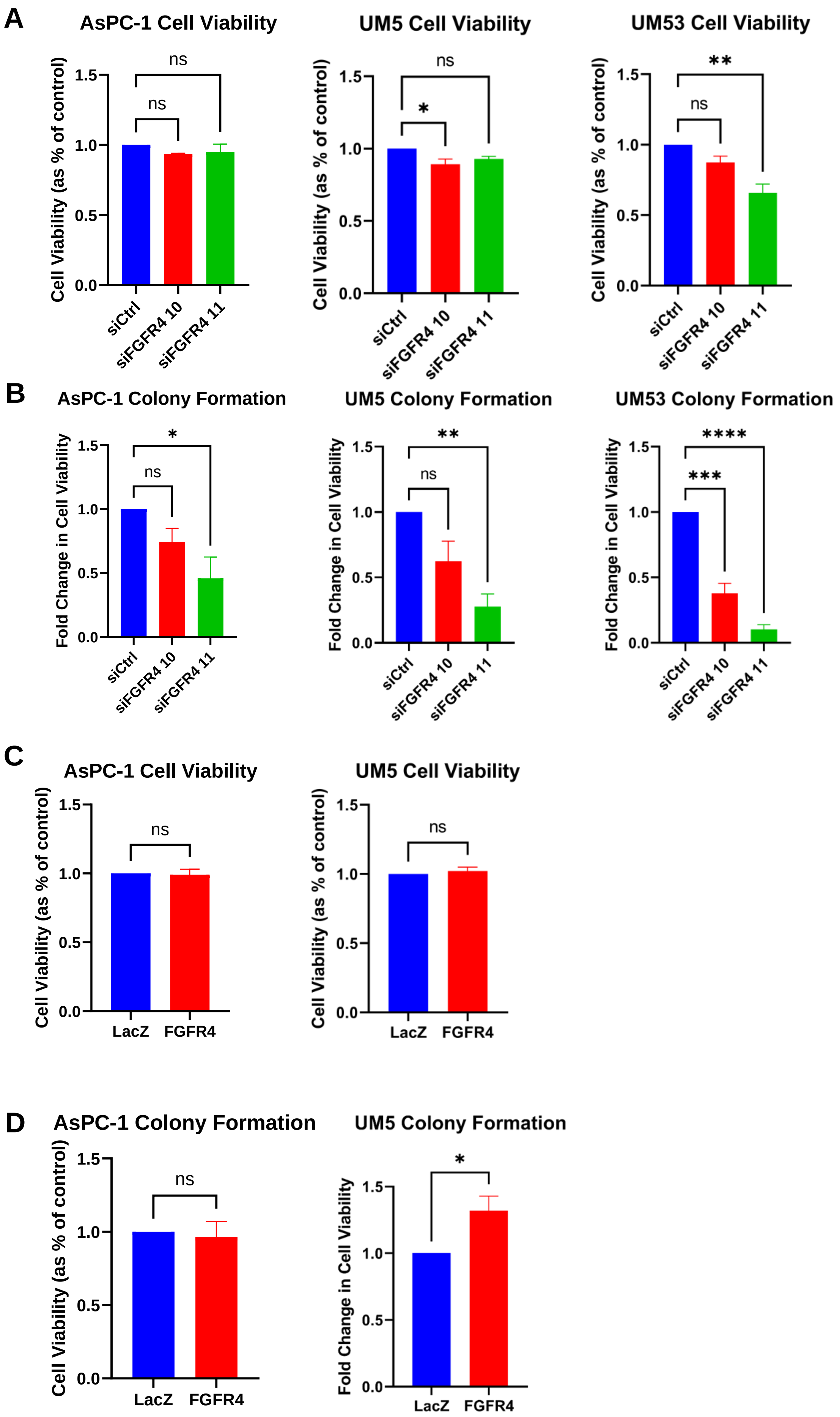

Supp Figure 4

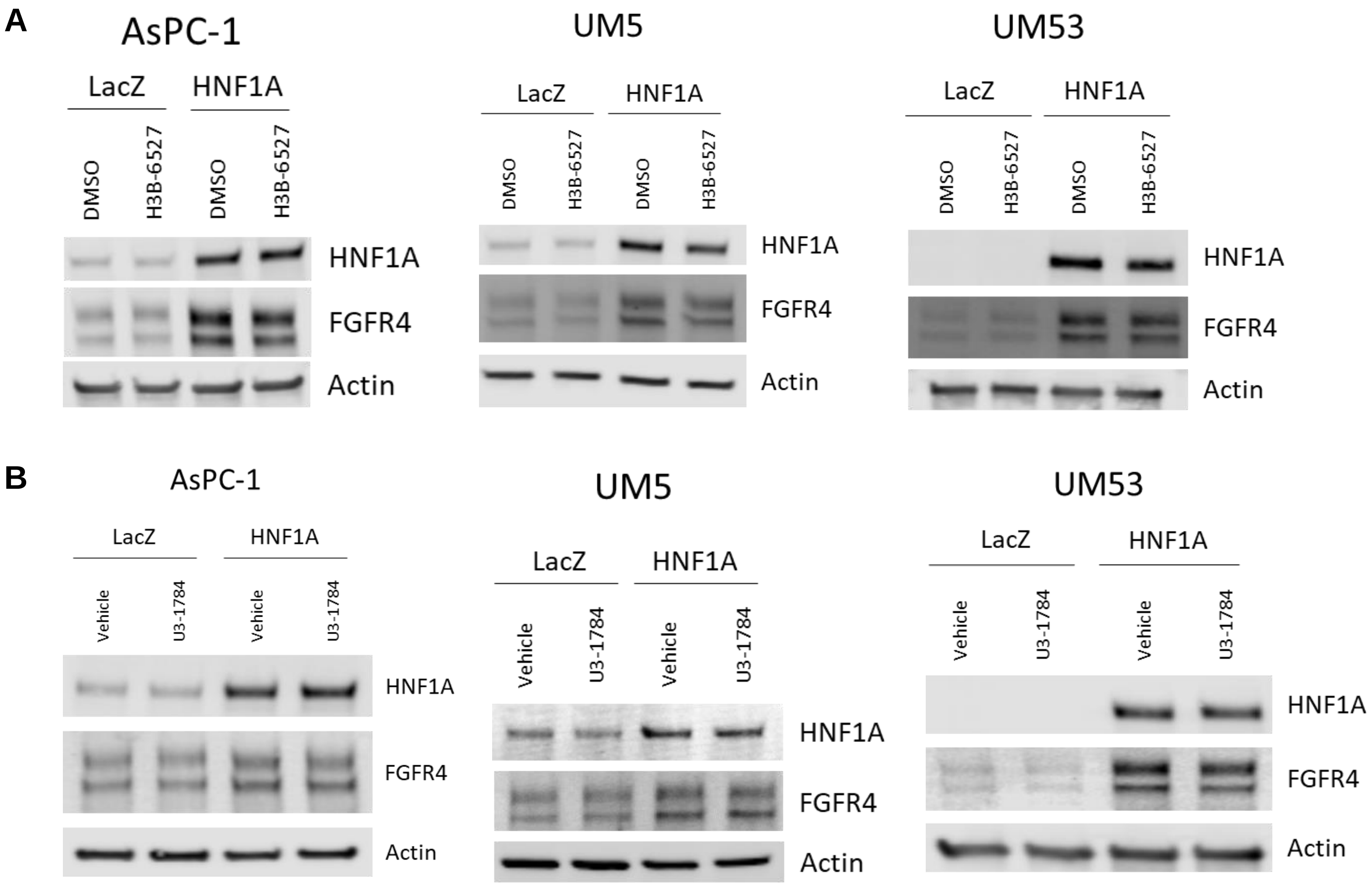

#### Supp Figure 5

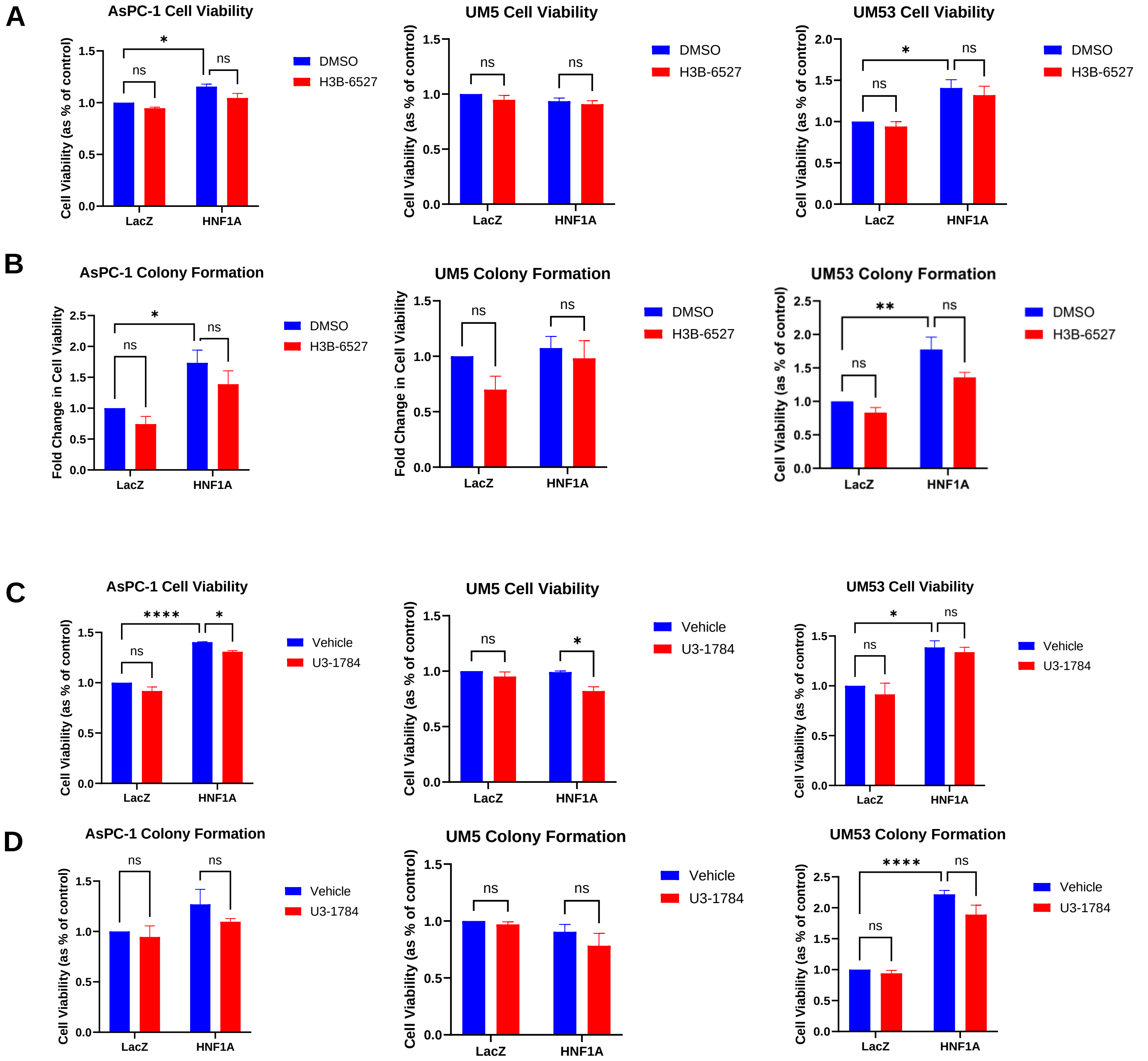
